## Supplementary Figure 1 for "Position representations of moving objects align with real-time position in the early visual response"

### 1 Supplementary Figure 1

Control analyses were conducted to ensure that EEG decoding results were not confounded by systematic microsaccades, either due to neural activity related to motor planning, movement-related EEG artefacts or shifts in retinotopy [1]. Firstly, the same decoding analysis that was applied to the EEG response to the static stimuli was applied to the x-y position of the eyes (measured using concurrent eye-tracking). It was not possible to decode the position of the stimulus from eye position at any timepoint between 0 and 350ms (stimulus onset to 100ms after stimulus offset, as used for the EEG analysis), as shown in Figure 1a. This is in line with Blom et al. [2] and Salti et al. [3]; the former paper found the location of a flashed wedge around a circle could not be decoded based on eye movements, and the latter found that eye position contained information about stimulus location only after 500ms. Additionally, Tse et al. [4] found that abrupt onsets did not elicit eye movements unless a saccade was already being planned, which was not the case in the present experiment. Secondly, a multi-sample test for equal median distributions [5] showed that the median eye angle, averaged from 50 to 350ms post flash onset, was not significantly different for any of the outermost corner stimulus position: $P(11) = 2.00$, $p = .849$. This can be seen in the eye angle histogram in Figure 1b. Both these analyses suggest that participants' eye positions did not systematically vary according to the stimulus location. Therefore, we can conclude that the classifiers trained on the EEG response to the static stimulus did not exploit eye movements to determine the stimulus position. Furthermore, it has been shown that, when viewing motion, microsaccades occur parallel to perceived motion direction [6;7]. This implies that any microsaccades elicited by the moving stimulus in the current experiment should be informative about motion direction but not stimulus position. Since the classifiers were trained on stationary stimuli, they could not exploit any motion direction information and therefore these microsaccades could not have influenced our results. This convergence of evidence from the eyetracking data and previous literature gives us confidence that the results of the main analysis can not be explained by eye movements.

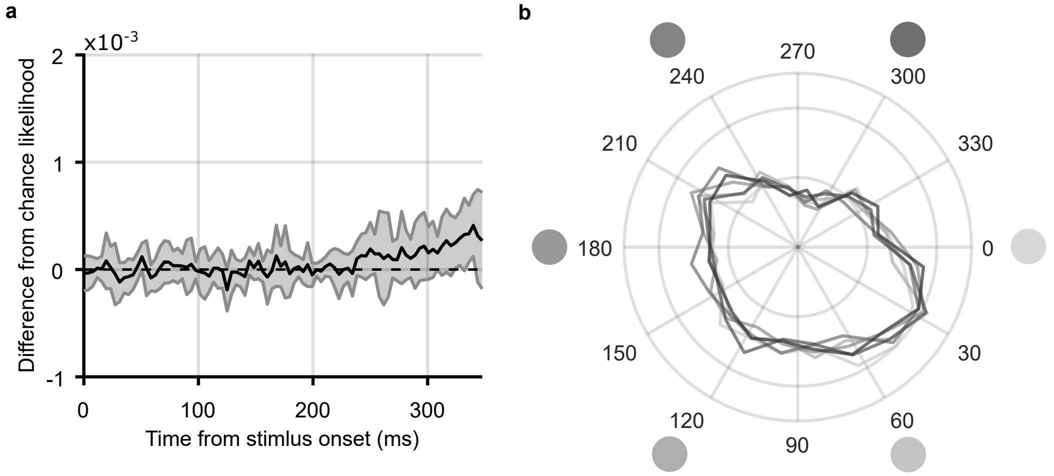


*Figure 1:* ***Analysis of eye movements. a)*** *Mean stimulus-position likelihood over time, calculated from classifiers trained to discriminate the location of the static stimulus using the x-y position of the eye at each timepoint following stimulus onset. This analysis parallels the analysis used on EEG data in Figure 2 (Results). Stimulus-position likelihood was not significantly above chance at any timepoint.* ***b)*** *Distribution of eye angles (density plot) following presentation of stimuli in the six outermost stimulus positions, 12.4dva from fixation. The angle of the stimulus around fixation is marked by the greyscale circles; corresponding coloured lines show the distribution of eye angle for that stimulus. For each trial and each participant, the average eye angle between 50ms and 350ms was included. It can be observed that the distribution of eye angle does not change depending on the angular position of the stimulus.*
